# Glycogen Synthase Kinase 3 induces multilineage maturation of human pluripotent stem cell-derived lung progenitors in 3D culture

**DOI:** 10.1101/410894

**Authors:** Ana Luisa Rodrigues Toste de Carvalho, Alexandros Strikoudis, Tiago J. Dantas, Ya-Wen Chen, Hsiao-Yun Liu, Richard B. Vallee, Jorge Correia-Pinto, Hans-Willem Snoeck

**Author notes:** Correspondence should be addressed to H.W.S.

## Abstract

Although strategies for directed differentiation of human pluripotent stem cells (hPSCs) into lung and airway have been established, terminal maturation of the cells remains a vexing problem. We show here that in Collagen I 3D cultures in the absence of glycogen synthase kinase 3 (GSK3) inhibition, hPSC-derived lung progenitors (LPs) undergo multilineage maturation into proximal cells arranged in pseudostratified epithelia, type I alveolar epithelial cells and morphologically mature type II cells. Enhanced cell cycling, one of the signaling outputs of GSK3 inhibition, plays a role in the maturation-inhibiting effect of GSK3 inhibition. Using this model, we show NOTCH signaling induced a distal at the expense of a proximal and ciliated cell fate, while WNT signaling promoted a proximal, club cell fate, thus implicating both signaling pathways in proximodistal specification in human lung development. These findings establish an approach to achieve multilineage maturation of lung and airway cells from hPSCs, demonstrate a pivotal role of GSK3 in the maturation of lung progenitors, and provide novel insight into proximodistal specification during human lung development.

## INTRODUCTION

The generation of lung and airway epithelial cells from human embryonic stem (ES) and induced pluripotent stem (iPS) cells, collectively termed human pluripotent stem cells (hPSCs), holds major promise for studies in cellular physiology, disease modeling and regenerative medicine. Directed differentiation attempts to recapitulate development. The respiratory system originates from buds on the ventral anterior foregut endoderm (AFE) that undergo branching morphogenesis culminating in the emergence of basal, goblet, club, ciliated and neuroendocrine cells in the stalks, which will become airways, while the distal tips widen into primitive alveoli (Swarr and Morrisey, 2015). Mature alveoli contain alveolar epithelial type I (ATI) cells, which are essential for gas exchange, and type II (ATII) cells, which produce surfactant, critical for the maintenance of alveolar integrity by reducing surface tension (Whitsett et al., 2010). Thus, to generate lung and airway epithelial cells from hPSCs definitive endoderm (DE), anterior foregut endoderm (AFE), ventral AFE and lung progenitors (LPs) are sequentially specified followed by further differentiation into a mixture of alveolar and airway cells (Chen et al., 2017; Dye et al., 2016; Dye et al., 2015; Firth et al., 2014; Gotoh et al., 2014; Huang et al., 2015; Huang et al., 2014; Jacob et al., 2017; Konishi et al., 2016; McCauley et al., 2017; Mou et al., 2012; Wong et al., 2012).

Several challenges remain however. Similar to other organs and tissues (Lancaster and Knoblich, 2014), terminal maturation remains elusive. We reported a 3D model consisting of lung bud organoids generated in suspension from early AFE followed by embedding in Matrigel (Chen et al., 2017), where branching morphogenesis with predominant generation of ATII cells ensued. These organoids were equivalent to the second trimester of human gestation. In other published models, maturation stage is not reported or is also equivalent to early fetal lung (**Suppl. Table 1**) (Dye et al., 2016; Dye et al., 2015; Firth et al., 2014; Gotoh et al., 2014; Huang et al., 2014; Jacob et al., 2017; Konishi et al., 2016; McCauley et al., 2017; Wong et al., 2012; Yamamoto et al., 2017). A method that is permissive for all mature lung epithelial cells would be useful to study lineage relationships, the mechanisms driving the specification of individual lineages, and the biology of mature human lung and airway cells, for example to investigate tropism of human respiratory viruses for specific cell types. A second challenge is that it is not known how proximodistal specification can be achieved *in vitro* while its mechanisms *in vivo* are unclear. Faithful proximodistal specification is a prerequisite to generate more homogeneous populations of specific epithelial cells. Recently it was reported that canonical Wnt agonism induced by the GSK3 inhibitor, CHIR9902 (CHIR), promoted specification of developmental lung progenitors (LPs) towards ATII cells, whereas withdrawal of CHIR induced a proximal fate. These studies used reporter lines to enrich for desired progenitor populations or identify desired differentiated lineages (Jacob et al., 2017; Longmire et al., 2012; McCauley et al., 2017), however, and are therefore not universally applicable. Several other reports also show the generation of ATII cells (Chen et al., 2017; Huang et al., 2014; Jacob et al., 2017; Yamamoto et al., 2017). Neither mature NGFR^+^ basal cells (BCs) (Rock et al., 2009), the stem cells of the airways, nor ATI cells were ever generated however, perhaps because both cell types arise late in development (Frank et al., 2016; Yang et al., 2018).

To address these issues, a culture model that does not rely on reporter lines and is permissive for the specification of all lung and airway lineages, thus allowing investigation of conditions that favor specific lineages, is required. Here we report a collagen I (Col I) 3D culture system that satisfies these criteria. We show that GSK3 inhibition, rather than favoring distal fates as reported previously (McCauley et al., 2017), promotes proliferation and inhibits differentiation, whereas withdrawal of GSK3 inhibition induces multilineage proximal and distal maturation, including of NGFR^+^ basal cells, morphologically mature ATII cells and cells with the morphology and marker expression of ATI cells. Furthermore, a WNT ligand could not recapitulate the effect of GSK3 inhibition, suggesting that this effect is not primarily mediated by canonical WNT signaling. Generic cell cycle inhibition, however, partially recapitulated the effect of CHIR withdrawal, suggesting a role for GSK3-mediated cell cycle regulation in maturation of LPs. We next used this model to show that after CHIR withdrawal NOTCH inhibition promotes proximal and inhibits distal development, thus identifying NOTCH signaling as one of the signaling pathways involved in proximodistal specification.

## Results

### Establishment of a 3D Col I model of human lung and airway lineage specification

Our published 2D culture protocol recapitulates development (Huang et al., 2015; Huang et al., 2014). For further studies, the 2D model posed two problems however. First, in large areas cell detachment occurred (Huang et al., 2015). Second, despite ample presence of cells expressing ATII markers, expression of the most specific ATII marker, SFTPC, was sporadic while ATI cells and BC-like cells were rare (Huang et al., 2015; Huang et al., 2014), and mature NGFR^+^ BCs absent. We therefore proceeded to culture in a 3D matrix.

We generated NKX2.1^+^FOXA2^+^ LPs, which lacked mature lung and mesenchymal markers, in 2D until day (d)25, when the purity of NKX2.1^+^FOXA2^+^ lung progenitors was maximal (90-98%), as described previously (Huang et al., 2015; Huang et al., 2014), and transferred these to Col I gels in the presence of factors used in 2D cultures (Huang et al., 2015; Huang et al., 2014) (CHIR, FGF10, KGF and dexamethasone, 8-bromo-cAMP and isobutylmethylxanthine (DCI) (Gonzales et al., 2002)) **(Fig. 1a, top).** The cells organized in strands enveloping empty lacunae **(Fig. 1a, upper left)** and almost uniformly expressed NKX2.1 (85.08±17.54%) **(Fig. 1a)**, FOXA2 (not shown) and the surface mucin, MUC1, whose apical expression indicated polarization **(Fig. 1a)**. Most cells also co-expressed variable amounts of SOX9 and SOX2 **(Fig. 1a)**, which in contrast to the mouse (Rockich et al., 2013), are commonly co-expressed in distal tips during human lung development(Chen et al., 2017; Nikolic et al., 2017). SFTPC^+^ cells were abundant, indicating that a 3D environment is critical for SFTPC expression **(Fig. 1a)**. The ATII marker, ABCA3, was detected as well **(Fig. 1a).** Some cells co-expressed ATI (PDPN) and ATII (SFTPB, SFTPC) markers **(Fig. 1a, arrows)** (Desai et al., 2014; Treutlein et al., 2014). Cells expressing markers of both ATI and ATII cells, called bipotential progenitors, are found at the distal tips of the developing lung and may differentiate into ATI or ATII cells (Desai et al., 2014; Treutlein et al., 2014), although more recent evidence suggests that the ATI-ATII fate decision is made earlier in development (Frank et al., 2016; Zacharias et al., 2018). Other ATI (CAV1, AKAP5, CLIC5, SCNN1A, Col IV, AQP5) (Treutlein et al., 2014) and proximal (CC10, MUC5B, P63,) markers were not observed **(Fig. S1)**, as were mesodermal (PDGFR*α*, CD90) and endothelial markers (CD31) (not shown).

**Figure 1.**
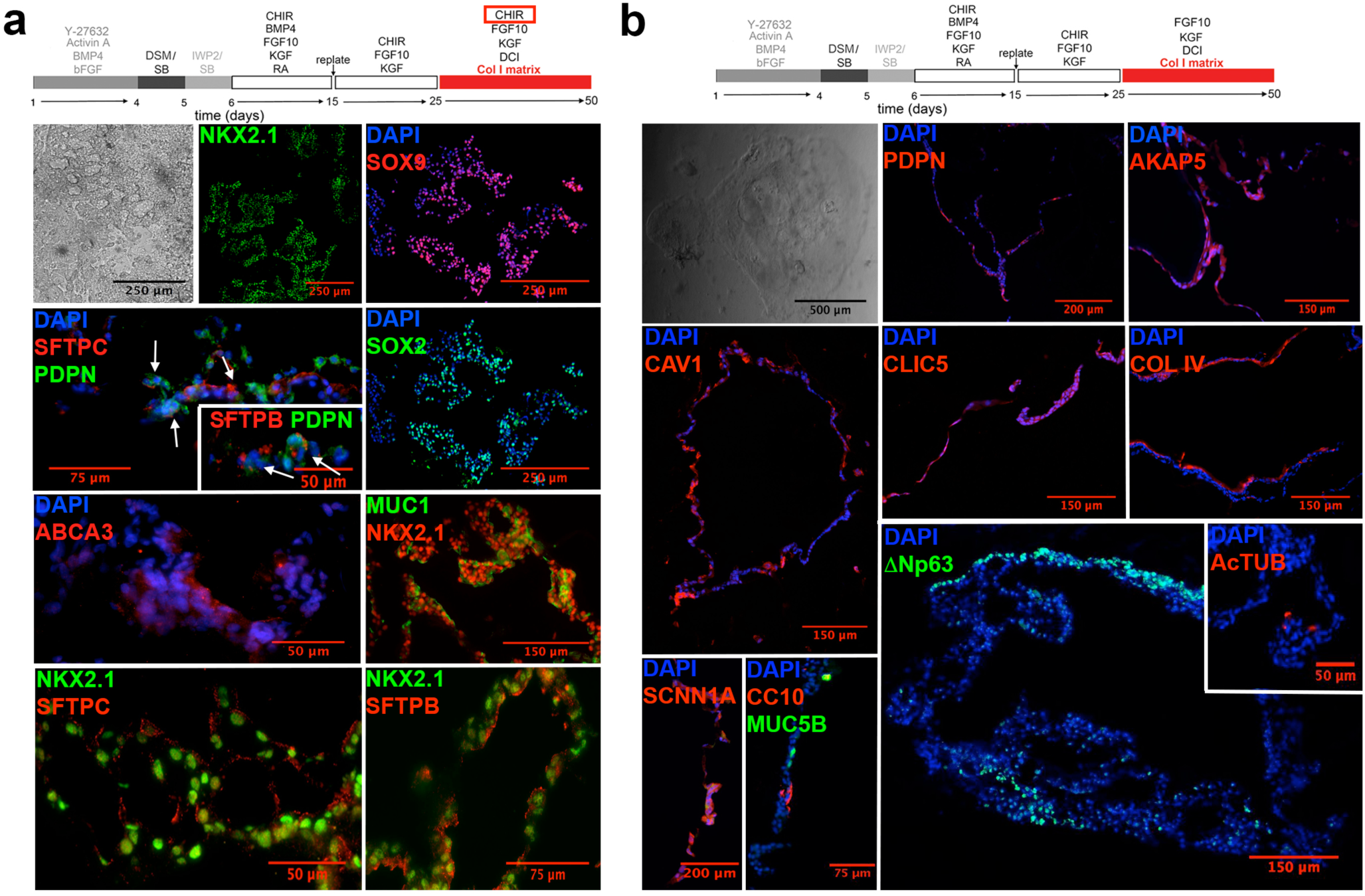
CHIR withdrawal induces maturation. **(a)** Brightfield and IF for indicated markers after culture of RUES2 ESCs in the presence of CHIR according to the protocol on top (representative of >12 experiments; arrows indicate cells co-expressing ATII and ATI markers). **(b)** Brightfield and IF for indicated markers after culture of RUES2 ESCs in the absence of CHIR according to the protocol on top (representative of >12 experiments).

Removing the GSK3 inhibitor at the initiation of the 3D cultures at d25 (CHIR−) **(Fig. 1b, top)** dramatically changed morphology and marker expression. Comparative IF for 18 markers is shown in **Fig. S1**. The cells were arranged in more discrete colonies, expressed the endodermal markers FOXA1, but mostly lost NKX2.1 **(Fig. S2a)**, which is later in lung development restricted to ATII and airway club cells. Markers for ATI cells, HOPX **(Fig. S2a)** and PDPN, and ATI markers distilled from the RNAseq data of Treutlein et al. (Treutlein et al., 2014) (COL IV, CAV1, CLIC5 and AKAP5) were widely expressed **(Fig. 1b, S1)** in flattened cells **(Fig. 1b, Fig. S1, S2a)**, some of which stained for SCNN1A, which is also expressed in ATI cells **(Fig. 1b, Fig. S1)** (Yang et al., 2016). COL IV was expressed basally **(Fig. 1b, Fig. S1)**, consistent with alveolar basement membrane production by ATI cells. These changes in morphology and combined marker expression indicate ATI specification. Other markers of mature ATI cells, such as AGER, were not detected however, suggesting still incomplete maturation. In addition, multiple clusters of P63^+^ **(Fig. 1b, Fig. S1, S2a)** and KRT5^+^ cells **(Fig. S1, S2a)**, a phenotype compatible with airway basal cells (BCs) (Rock et al., 2009), and cells expressing markers of differentiated airway cells (AcTUB^+^ (ciliated), CC10^+^ (club) and MUC5B^+^ (goblet)) were observed **(Fig. 1b, Fig. S1)**. These expression patterns suggest that Col I culture of LPs in the absence of CHIR induced multilineage, proximal and distal differentiation. Light sheet microscopy **(Fig. 2a)** showed irregularly shaped structures with markers for airway cells (MUC5B, AcTUB, CC10) occurring in discrete peripheral clusters, the BC marker KRT5 lining the outside of the structures, and the distal markers HT2-280 (ATII cells) and CAV1 (ATI cells) expressed more diffusely. Although Col I cultures do not generate structurally appropriate lung organoids, these qualitative findings confirm that CHIR withdrawal induces multilineage differentiation and maturation.

**Figure 2.**
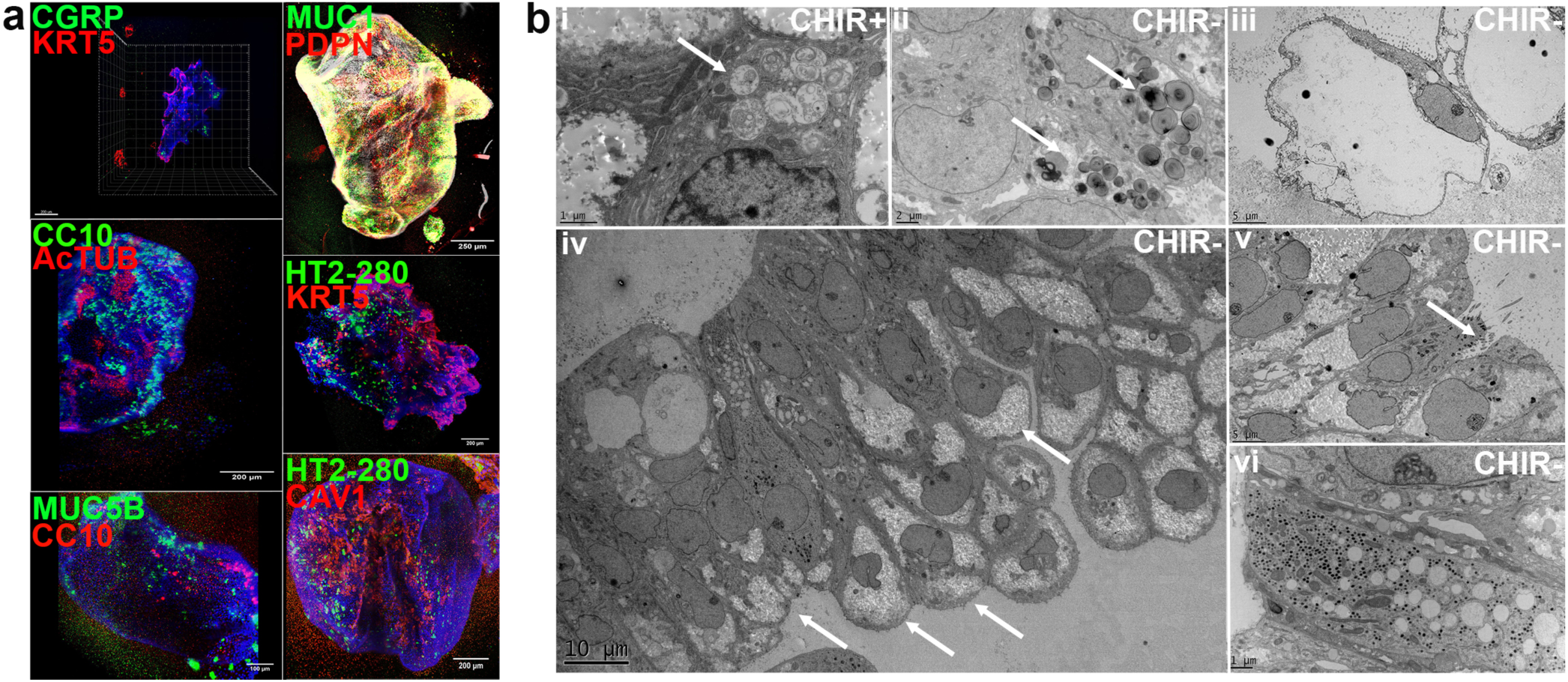
Light sheet and transmission electron microscopy of CHIR− cultures. **(a)** Whole mount IF of the indicated differentiation markers in CHIR− cultures at d50 (representative of 3 independent experiments). **(b)** Transmission electron microscopy of CHIR+ and CHIR− cultures at d50; **(i)** cell with immature LBs and multivesicular bodies (arrow) in CHIR+ conditions; **(ii)** cell with mature LBs in CHIR− conditions (arrows); **(iii)** Flat cells compatible with ATI cells in CHIR− conditions; **(iv)** Pseudostratified epithelium in CHIR− conditions, arrows indicate droplet-shaped cells; **(v)** Multiciliated cell in CHIR− conditions (arrow); **(vi)** Club cell in CHIR− conditions (all data from i-vi representative of n=1).

### Electron microscopy indicates enhanced multilineage maturation in CHIR− cultures

Transmission electron microscopy (TEM) of CHIR+ cultures showed mostly cuboidal cells **(Fig. 2b(i))** with apical cytoplasmic projections **(Fig. S2b)**, suggestive of polarization. Many contained lamellar bodies (LBs) at various stages of maturation, though most appeared as multivesicular bodies, the precursors of LBs (Weaver et al., 2002) **(Fig. 2b(i), S2b, arrows)**. In CHIR− conditions, cells with LBs were much sparser, consistent with reduced expression of ATII markers, but contained uniformly mature, electron-dense LBs **(Fig. 2b(ii), S2c, arrows)**, indicating maturation. Elongated cells **(Fig. 2b(iii), S2d)** lining a lumen that appeared free of matrix and with, in some instances, electron-dense material compatible with basement membrane **(Fig. S2e, arrow)**, a morphology compatible with ATI cells, were present. We observed multiple areas with a pseudostratified epithelium **(Fig. 2b(iv),(v))**, consisting of basal droplet-shaped cells, a morphology consistent with BCs **(Fig. 2b(iv) arrows)**, and club cells with secretory granules **(Fig. 2b(iv,vi))** interspersed with more rare ciliated cells **(Fig. 2b(v), S2f, arrow)** reaching the lumen. Other luminal cells had indeterminate morphology and may be precursors of goblet or club cells. These morphological findings are consistent with the immunofluorescence data, and indicate that withdrawal of CHIR induced multilineage maturation.

### Quantitative analysis of CHIR+ and CHIR− Col I cultures

To further substantiate the induction of multilineage differentiation in the absence of CHIR, we quantified differentiation in the cultures. In CHIR− cultures cell density was significantly lower than in CHIR+ cultures **(Fig. 3a,b)**, 15±7×10^6^ vs 163±45×10^6^ cells/10^6^ ESCs (n=4), indicating that CHIR supports proliferation of progenitors. mRNAs encoding markers for ATI, ciliated, club, neuroendocrine and basal cells were massively upregulated (10^2^ to 10^6^-fold) in CHIR− compared to CHIR+ cultures **(Fig. 3c)**. Although by TEM ATII cells in CHIR− cultures were morphologically mature, mRNA for ATII markers were down regulated in CHIR− compared to CHIR+ conditions however, most likely because cells expressing ATII markers are precursors for both mature ATII and ATI cells (Nabhan et al., 2018; Zacharias et al., 2018) and are the most abundant in the CHIR+ condition. These data therefore support the conclusion that CHIR withdrawal induced multilineage maturation. Quantification of IF images confirmed a decreased fraction of NKX2.1^+^ nuclei (24.16±18.59%) **(Fig. 3d)** and increased proportion of P63^+^ nuclei (34.52±2.25%) **(Fig. 3e)**. Furthermore, quantification of non-nuclear markers by measuring fluorescent area relative to DAPI area showed an increase in all tested differentiation markers in CHIR− compared to CHIR+ cultures **(Fig. 3f)**. Flow cytometry revealed an increase in the proportion of cells expressing HT2-280 (4.89±1.06% vs 0.75±0.28%) **(Fig. 4a)**, a marker for more mature ATII cells, and a strikingly increased fraction and absolute number of ITGA6^+^ITGB4^+^ cells, which were undetectable in CHIR+ conditions, but reached 24±12% in CHIR− conditions **(Fig. 4b)**. This phenotype is associated with airway progenitors(Rock et al., 2009). To further confirm the BC-like identity of these cells, we purified ITGA6^+^IGTB4^+^ cells using fluorescence-activated cell sorting and stained for BC markers. All cells co-expressed the BC markers p63, MUC1, KRT5, SOX2, and PDPN **(Fig. 4c)**, indicating that these are indeed BC-like. Furthermore, in human adult lung, ITGB4 was expressed specifically in the basal layer of airway epithelium, while ITGA6 was also sporadically expressed distally **(Fig. 4d)**. Finally, after longer culture, the adult BC marker, NGFR (Huang et al., 2014; Rock et al., 2009) was amply detected in epithelial structures in CHIR− conditions **(Fig. 4e)** with EPCAM^+^NGFR^+^ cells comprising 16.5±5.85% (4.56±1.8×10^4^ cells cm^−2^) of the total number of cells at d80 **(Fig. 4f)**. We next isolated ITGB4^+^NGFR^+^ cells, and expanded these on 3T3-J2 feeders in the presence of the ROCK inhibitor, Y29632(Butler et al., 2016). After 2 passages, during which the cells maintained their ITGB4^+^NGFR^+^ phenotype **(Fig. 4g)** and displayed a doubling time of approximately 0.8 days **(Fig. 4h)**, the cells were plated in air-liquid interphase cultures, where they differentiate into goblet, club and ciliated cells after 6 weeks **(Fig. 4i).** These observations demonstrate that mature NGFR^+^ BCs were generated. Taken together, quantitative analysis of the cultures indicates induction of multilineage differentiation and maturation in the absence of CHIR.

**Figure 3.**
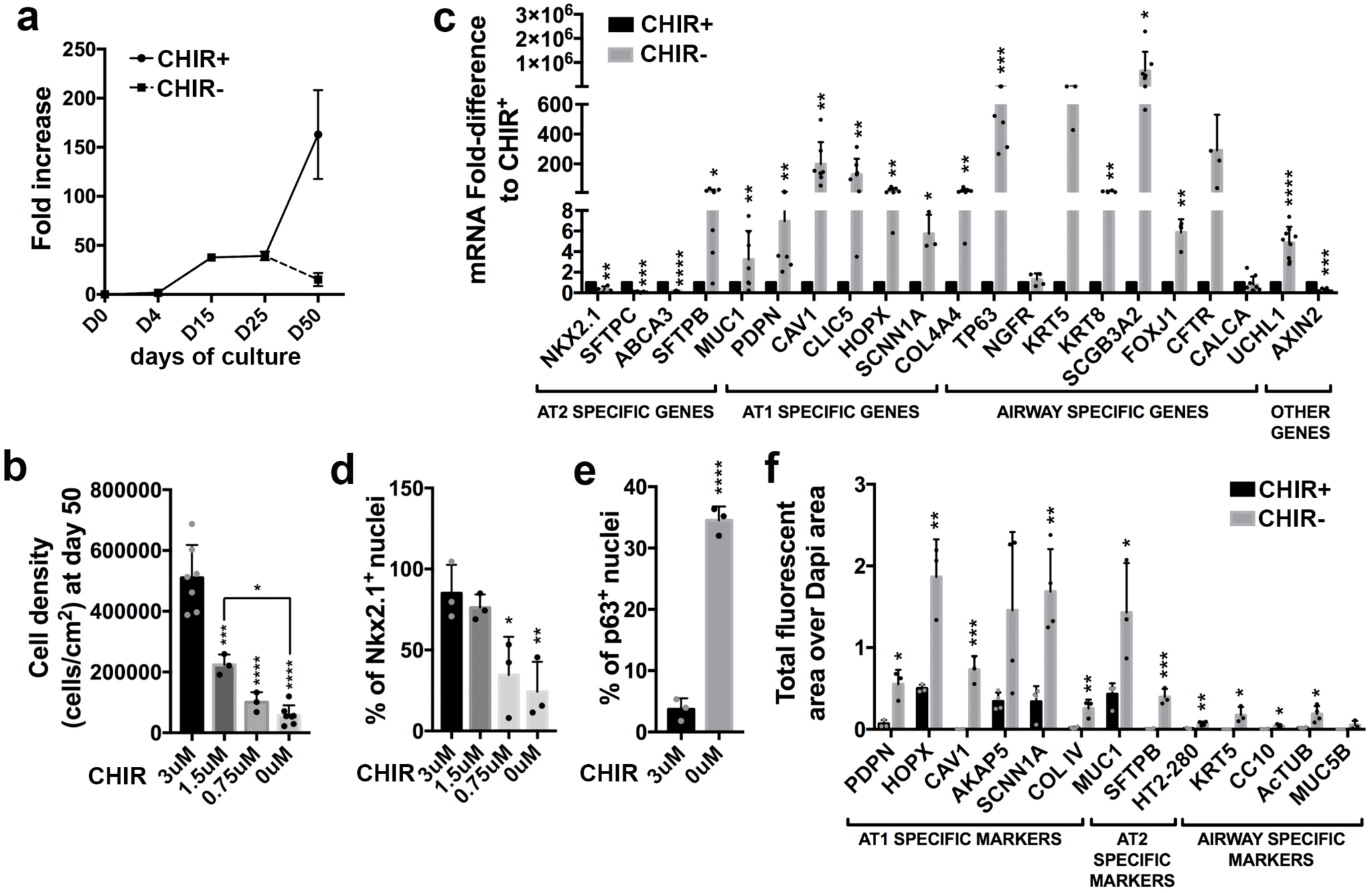
Quantitative analysis of CHIR+ and CHIR− cultures. **(a)** Cellular expansion starting from undifferentiated RUES2 ESCs in CHIR+ and CHIR− cultures (mean± s.d., n=4 independent experiments). **(b)** Cellular expansion from d25 to d50 in the presence of varying concentrations of CHIR (mean± s.d., 3μM and 0 μM n=7; 1.5 μM and 0.75 μM n=3, *p<0.05, ***p<0.001, ****p<0.0001). **(c)** Expression of mRNA for differentiation markers in CHIR− cultures relative to CHIR+ cultures (mean± s.d., n=3 to 7 independent experiments, *p<0.05, **p<0.01, ***p<0.001, ****p<0.0001). **(d)** Fraction of NKX2.1^+^ nuclei in d50 cultures after culture from d25 to d50 in varying concentrations of CHIR (mean± s.d., n=3, * p<0.05, **p<0.01). **(e)** Fraction of p63^+^ nuclei in d50 cultures after culture from d25 to d50 in varying concentrations of CHIR (mean± s.d., n=3 independent experiments, p<0.0001). **(f)** Relative quantification of IF for non-nuclear protein as determined by the ratio between fluorescent area for a given marker and nuclear DAPI area (mean± s.d., n=4 independent experiments, *p<0.05, **p<0.01, ***p<0.001).

**Figure 4.**
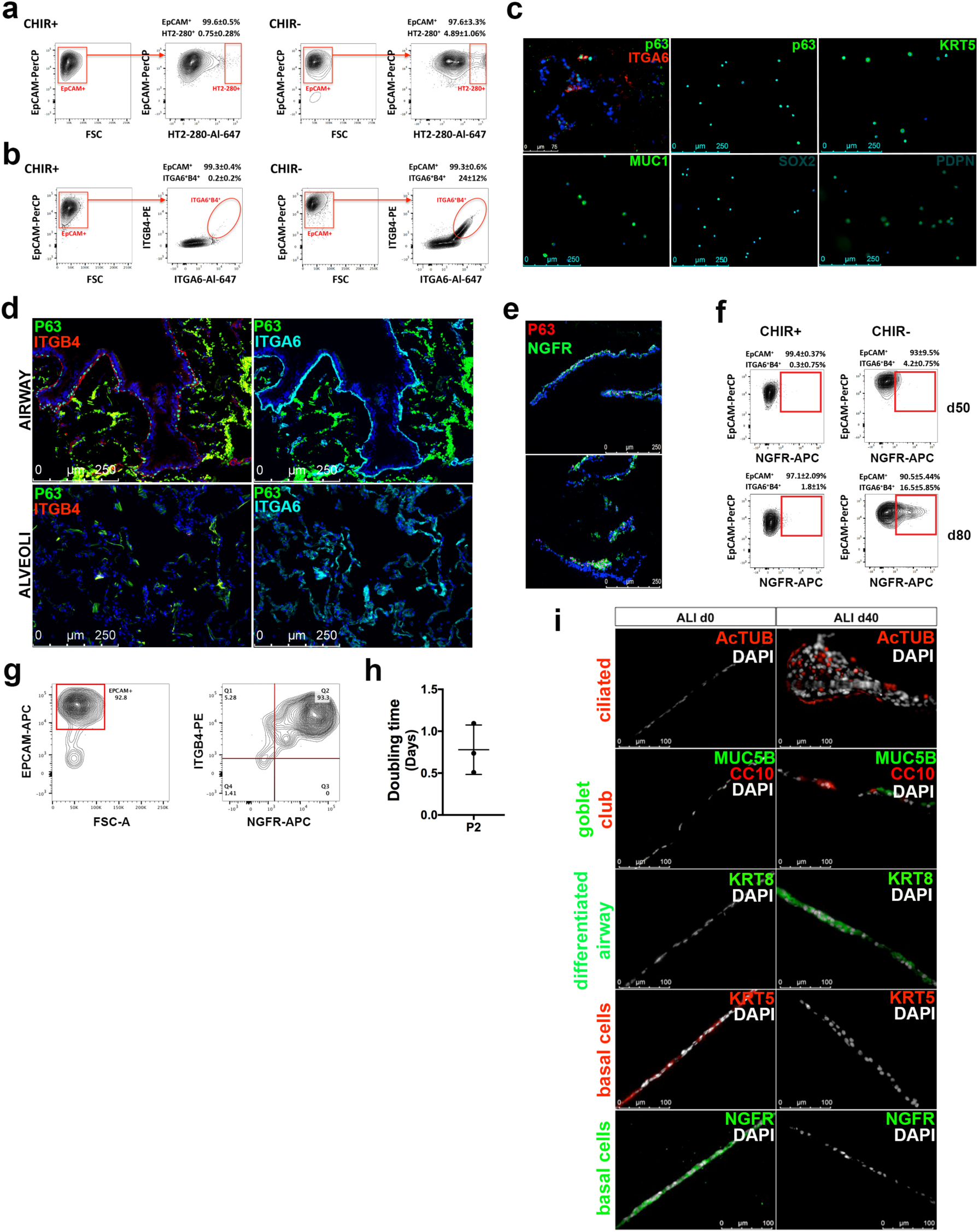
Airway progenitors and distal cells in CHIR− cultures. **(a)** Representative example and statistical analysis of the flow cytometric profile after staining for EPCAM and HT2-280 of cells from CHIR+ and CHIR− cultures (n=4 independent experiments). **(b)** Representative example and statistical analysis of the flow cytometric profile after staining for EPCAM, ITGA6 and ITGB4 of cells from CHIR+ and CHIR− cultures (n=4 independent experiments). **(c)** Staining of CHIR− cultures for p63 and ITGA6 (upper left corner) and IF for the indicated markers on preparations of ITGA6^+^ITGB4^+^ cells isolated from CHIR− cultures by flow cytometric cell sorting (representative of 1 experiment). **(d)** Expression patterns of ITGB4, ITGA6 and p63 in human adult airway and alveoli by IF staining (representative of 1 human adult lung sample). **(e)** IF staining for NGFR and p63 in cultures differentiated in CHIR− conditions until d120 (representative of n=3 independent experiments). **(f)** Representative example and statistical analysis of the flow cytometric profile after staining for EPCAM and NGFR of cells from CHIR+ and CHIR− cultures at d50 and d80 of the differentiation protocol (n=6 independent experiments). **(g)** Representative analysis of expression of EPCAM, ITGB4 and NGFR after two passages of purified NGFR^+^ITGB4^+^ cells from d80 CHIR− cultures in the presence of 3T3-J2 cells and RI. **(h)** Doubling time of cells in (g) (n=3). **(i)** Immunofluorescence of cells in (g) at initiation and after 40 days of air-liquid interphase culture.

### Genome-wide expression analysis confirms multilineage maturation in CHIR− Col I cultures

Genome-wide expression analysis and cross-referencing with the single cell expression data of Treutlein et al. in fetal mouse lung (Treutlein et al., 2014) revealed that although many transcripts associated with the ATII lineage were variably affected **(Fig. 5a)**, most transcripts indicative of ATI, ciliated and club cell differentiation as well as BC markers and secreted mucins **(Fig. 5b-f)** were upregulated, confirming multilineage maturation in CHIR− cultures.

**Figure 5.**
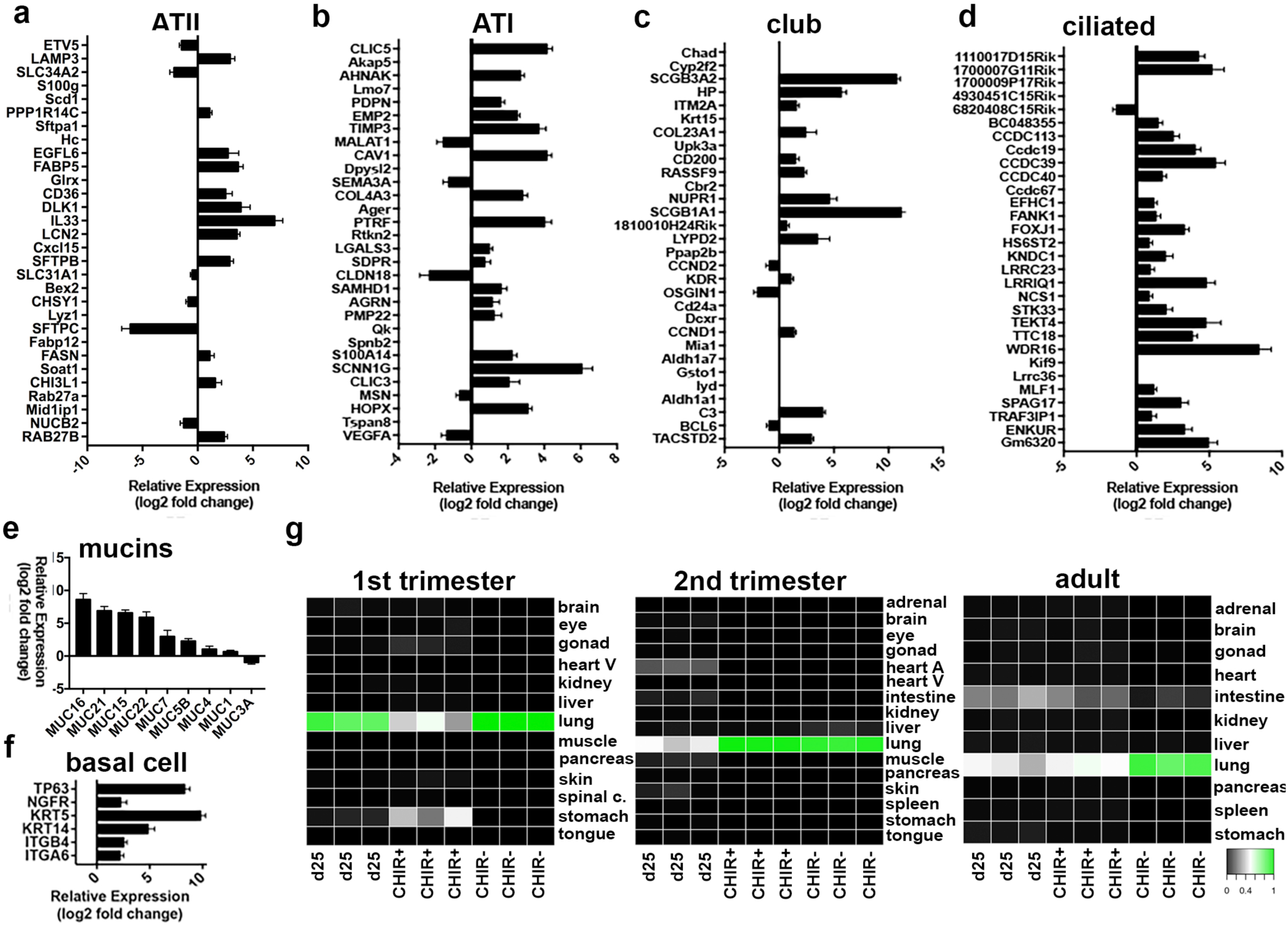
Genome-expression analysis. **(a-d)** Genes significantly differently expressed by RNAseq in CHIR− compared to CHIR+ cultures classified as marking ATII, ATI, ciliated and club cell markers according to Treutlein et al. Data are only shown for these genes classified by Treutlein et al. that were significantly different between CHIR− and CHIR+ conditions. **(e)** Mucin genes significantly differently expressed by RNAseq in CHIR− compared to CHIR+ cultures. **(f)** Genes expressed in BCs significantly differently expressed by RNAseq in CHIR− compared to CHIR+ cultures (all data from a-f mean± s.d., n=3 independent experiments, all data shown FDR<0.05). **(g)** Comparison of genome-wide expression of LPs (d25), and d50 CHIR+ and CHIR− cultures with the KeyGenes database (n=3 independent experiments).

To assess maturity with respect to human development, we cross-referenced the RNAseq data with the KeyGenes database, which contains expression profiles of human organs during 1^st^ and 2^nd^ trimesters of gestation and adulthood(Roost et al., 2015). D25 LPs showed the best match with the first trimester lung, consistent with their early stage of development. Cells from CHIR+ cultures corresponded to 2^nd^ trimester. Cells from CHIR− cultures corresponded to 1^st^ and 2^nd^ trimester and to adult lung **(Fig. 5g)**, a finding most likely explained by the large diversity of cell types at varying stages of differentiation. The fact that only cells cultured in CHIR− matched with human adult lung however is consistent with further maturation in this condition.

Finally, single cell RNAseq and t-SNE analysis showed several clusters of cells in CHIR− conditions that could be assigned to specific cell types (airway progenitors, club cells, ATI cells and NE cells) based on key markers **(Fig. S3)**. Using the same markers, none of these populations could be identified in CHIR+ conditions **(Fig. S3)**. These data again demonstrate multilineage differentiation induced by CHIR removal.

### Effect of varying culture conditions

We next compared Col I and Matrigel as 3D media. Although a differentiation-inducing effect was also noted in CHIR− conditions in Matrigel **(Fig. S4a,b)**, expression of markers of mature cells was significantly lower than in Col I **(Fig. S4c)**. Col I is therefore more permissive for multilineage maturation than Matrigel. It has also been suggested that replacing KGF by FGF2 and higher concentrations of FGF10 are more efficient for generating proximal cells (McCauley et al., 2017). However, except for SFTPC, expression of most markers was lower in this condition **(Fig. S4d)**, although proximal and distal markers could be detected by IF **(Fig. S4e)**.

Finally, as CHIR is an agonist of canonical WNT signaling, which shows stage-specific effects during lung development (Swarr and Morrisey, 2015), we explored early (d15) and late (d35) CHIR withdrawal. Delayed withdrawal at d35 increased cell density **(Fig. S5a)**, and even further increased the absolute number of both ITGA6^+^IGTB4^+^ BC-like cells (5.42±1.6×10^4^ cells cm^−2^) **(Fig. S5b)** and HT2-280^+^ ATII cells (4.59±3×10^4^ cells cm^−2^) **(Fig. S5c)**, as well as mRNA expression for most differentiation markers **(Fig. S5d)** compared to CHIR withdrawal at d25. IF confirmed the ample presence of differentiation markers when CHIR was withdrawn at d35 **(Fig. S5e)** and quantification showed increased expression of all markers at the protein level **(Fig. S5f)**. This finding is consistent with additional CHIR− driven progenitor expansion followed by multilineage differentiation after CHIR withdrawal. In contrast, early CHIR withdrawal at d15 severely compromised the competence of LPs to express differentiation markers **(Fig. S5g)**.

Taken together, Col I is optimally permissive for multilineage differentiation, while timing of CHIR withdrawal is critical to assure competence of LPs to undergo multilineage differentiation, but does not affect proximodistal specification.

### The effect of CHIR is not reproduced by canonical WNT signaling

CHIR is frequently used as an agonist of canonical WNT signaling(Jacob et al., 2017; McCauley et al., 2017), as canonical WNT signaling inhibits GSK3 Ser/Thr kinase activity, thereby stabilizing *β*-catenin (Willert and Nusse, 2012). WNT signaling. CHIR increased AXIN2 mRNA (**Fig. 3c)**, nuclear *β*-catenin and inhibitory S9 phosphorylation of GSK3 **(Fig. 6a)**, indicating GSK3 inhibition and WNT activation, as expected. We therefore assessed the specific effect of the canonical WNT ligand, WNT3A, in CHIR− conditions. WNT3A did not affect cellular expansion **(Fig. 6b)** and even further increased the fraction of ITGA6^+^ITGB4^+^ cells (31.8±12.95%) **(Fig. 6c)**, the number of P63^+^ nuclei (51.5±9.78%) **(Fig. 6d)** and KRT5 pixel number (**Fig. 6e)**, while not affecting the fraction of HT2-280^+^ ATII cells (not shown) (comparative IF for 18 markers shown in **Fig. S1**). WNT3A did not increase TP63 or KRT5 mRNA however, but increased mRNAs for club cell markers (NKX2.1, SFTPB and SCGB3A2) **(Fig. 6f)**. WNT3A therefore neither qualitatively nor quantitatively recapitulated the effect of CHIR, and contrary to previous reports (Jacob et al., 2017; McCauley et al., 2017), our findings indicate that WNT signaling does not promote distal at the expense of proximal fate, but in fact has an opposite effect by modestly favoring a proximal and club cell fate. Because WNT3A neither quantitatively nor qualitatively replicated the effects of CHIR, it is unlikely that the differentiation-inhibiting effects of CHIR depend on WNT agonism.

**Figure 6.**
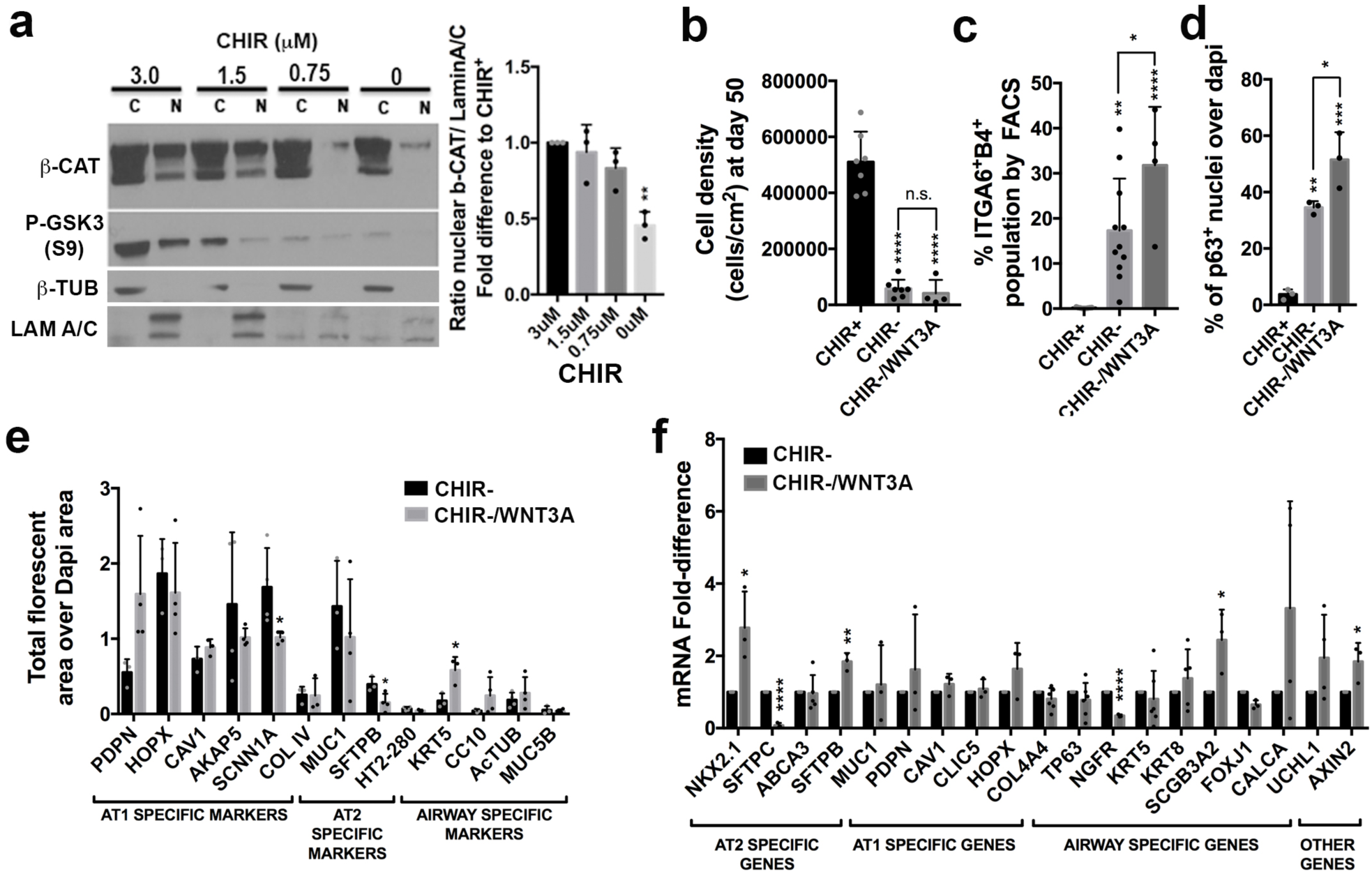
Role WNT signaling in the effect of CHIR. **(a)** WB on nuclear and cytoplasmic fractions for *β*-catenin and S9 phospho-GSK3 in cultures treated with varying concentration of CHIR (representative of 3 experiments) and quantification of WB on nuclear fractions of cultures treated with varying concentration of CHIR for nuclear b-catenin (mean± s.d., n=3 independent experiments, **p<0.01). **(b)** Cell density in CHIR+, CHIR− and CHIR− cultures treated with 100ng/ml WNT3A (mean± s.d., CHIR+ and CHIR− n=7; CHIR− /WNT3A n=4 independent experiments, ****p<0.0001). **(c)** Fraction of ITGA6^+^ITGB4^+^ cells in CHIR+, CHIR−, and CHIR− cultures treated with 100ng/ml WNT3A as determined by FCM. (mean± s.d., n between 4 to 11 independent experiments, *p<0.05, **p<0.01, ****p<0.0001). **(d)** Fraction of p63^+^ cells in CHIR+, CHIR− and CHIR− cultures treated with 100ng/ml WNT3A as determined by IF (mean± s.d., n= 3 independent experiments, *p<0.05, **p<0.01, ****p<0.0001). **(e)** Relative quantification of IF for non-nuclear protein as determined by the ratio between fluorescent area for a given marker and nuclear DAPI area in CHIR− cultures and CHIR− cultures treated with 100ng/ml WNT3A (mean± s.d., n=4 independent experiments, *p<0.05). **(f)** Expression of mRNA for differentiation markers in CHIR− cultures treated with 100ng/ml WNT3A relative to CHIR− (mean± s.d., n between 3 to 6 independent experiments, *p<0.05, **p<0.01, ****p<0.0001).

### The effect of cell cycle regulation by GSK3

GSK3 integrates inputs from multiple signaling pathways and one of its outputs is cell cycle regulation. As in neural progenitors, cell cycle arrest induces differentiation (Calegari et al., 2005; Calegari and Huttner, 2003; Kim et al., 2009; Lange et al., 2009; Roccio et al., 2013), we tested the hypothesis that some of the effects of CHIR withdrawal might be due to a lengthening of the cell cycle, while GSK3 inhibition might prevent differentiation by enhancing cycling and in doing so prevent maturation. We added two cell cycle inhibitors that act through distinct mechanisms, a CDK4/6-inhibitor (PD0332991, PD) (Dickson, 2014) and a CDC7 inhibitor (XL413, XL), to CHIR+ cultures. Although subtle differences were observed between both inhibitors, both PD and XL decreased cellularity **(Fig 7a,d)**, upregulated mRNAs for all differentiation markers and strikingly increased expression of mRNAs for distal and ATII markers **(Fig. 7b,e)**. IF confirmed widespread induction of distal markers in CHIR+ cultures in the presence of PD or XL **(Fig. 7c,f,** comparative IF for 18 markers shown in **Fig. S6)**. Furthermore, in the presence of either PD or XL the frequency of the ATII marker, HT2-280, which was virtually absent in CHIR+ conditions, was similar to that observed in CHIR− conditions **(Fig. 7g,h)**. On the other hand, as analyzed by FCM, induction of ITGA6 and ITGB4 by PD and XL **(Fig. 7j)** was weak compared to CHIR withdrawal **(Fig. 7i)**. IF and flow cytometry are therefore entirely consistent with mRNA expression data. To morphologically assess maturation in CHIR+ cultures in the presence of cell cycle inhibitors, we performed TEM of CHIR+/PD and CHIR+/XL cultures. These studies revealed predominant presence of cells with LBs. However, these were not mature, and most appeared as multivesicular bodies **(Fig. 7j)**. Similar to CHIR withdrawal, cell cycle inhibition therefore induced differentiation, but, in contrast to CHIR withdrawal, strongly favored distal cells and in particular the ATII lineage without promoting full maturation. We conclude that, cell cycle lengthening plays a role in the differentiation-inducing effect of CHIR withdrawal, but does not fully explain its effect on multilineage differentiation and maturation.

**Figure 7.**
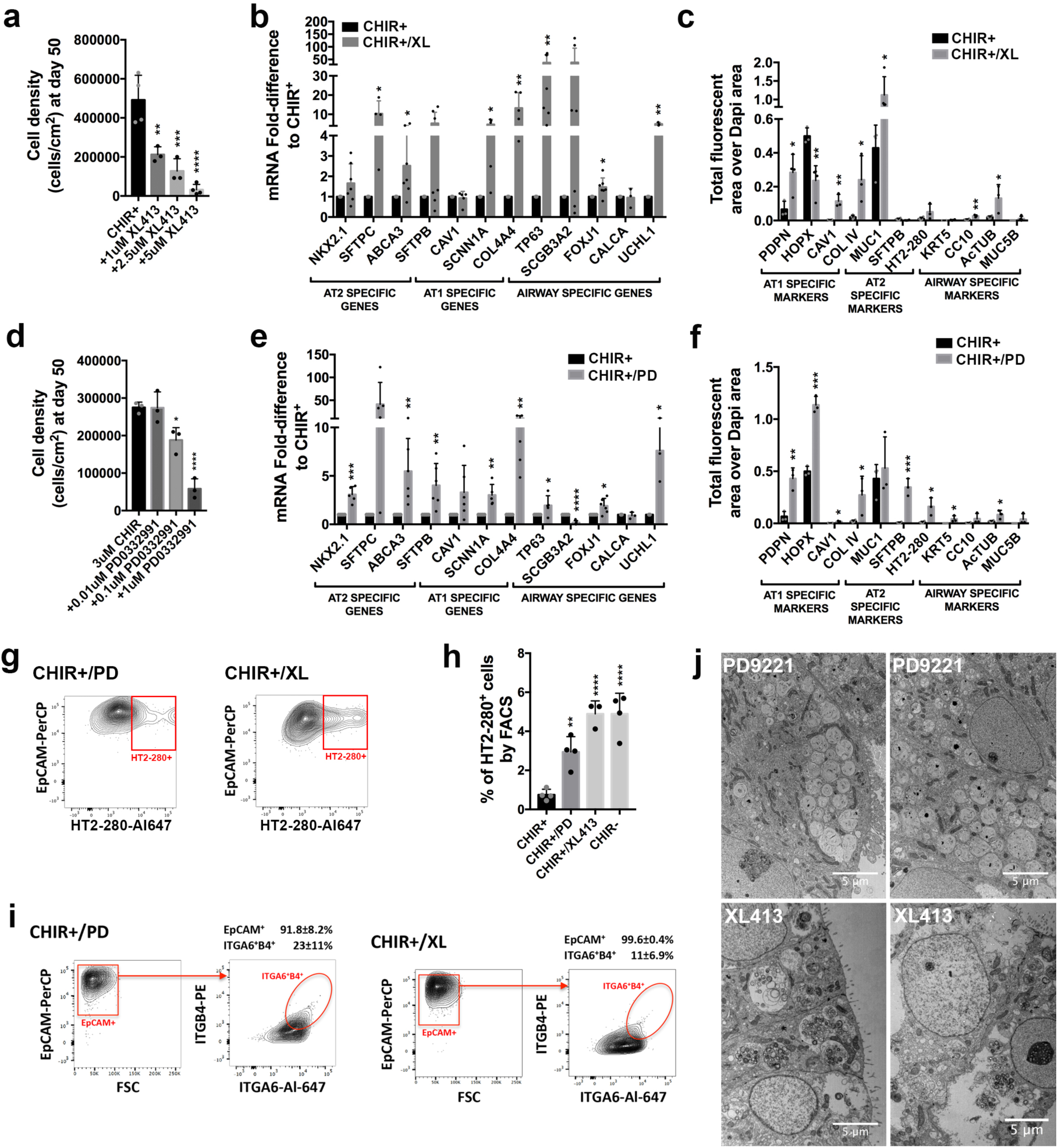
Effect of cell cycle inhibition on lung progenitor maturation. **(a)** Cell density in CHIR+ cultures supplemented with increasing concentrations of the CDC7 inhibitor XL413 (mean± s.d., n=3 independent experiments, **p<0.01, ***p<0.001, ****p<0.0001). **(b)** Expression of mRNA for differentiation markers in CHIR+ cultures treated with 5μM of XL413 relative to CHIR+ cultures (mean± s.d., n between 3 to 7 independent experiments, *p<0.05, **p<0.01, ***p<0.001, ****p<0.0001). **(c)** Relative quantification of IF for non-nuclear protein as determined by the ratio between fluorescent area for a given marker and nuclear DAPI area in CHIR+ cultures and CHIR+ cultures treated with treated with 5μM of XL413 (mean± s.d., n=3-4 independent experiments, *p<0.05, **p<0.01). **(d)** Cell density in CHIR+ cultures supplemented with increasing concentrations of the CDK4/6 inhibitor PD0339221 (mean± s.d., n=3 independent experiments, *p<0.05, ****p<0.0001). **(e)** Expression of mRNA for differentiation markers in CHIR+ cultures treated with 1μM of PD0332991 relative to CHIR+ cultures (mean± s.d., n between 3 to 6 independent experiments, *p<0.05, **p<0.01, ***p<0.001, ****p<0.0001). **(f)** Relative quantification of IF for non-nuclear protein as determined by the ratio between fluorescent area for a given marker and nuclear DAPI area in CHIR+ cultures and CHIR+ cultures treated 1μM of PD0332991 (mean± s.d., n=3-4 independent experiments, *p<0.05, **p<0.01, ***p<0.001). **(g)** Representative example of the flow cytometric profile after staining for EPCAM and HT2-280 of cells from CHIR+ cultures in the presence of either PD0332991 or XL413 (n=4). **(h)** Frequency of HT2-280^+^ cells in CHIR+, CHIR− and CHIR+ cultures supplemented with either PD0332991 or XL413 (mean± s.d., n= 3-4 independent experiments, **p<0.01, ****p<0.0001 compared to CHIR+). **(i)** Representative example and statistical analysis of the flow cytometric profile after staining for EPCAM, ITGA6 and ITGB4 of cells from CHIR+ cultures in the presence of either PD0332991 or XL413 hydrochloride (n=4). **(j)** Representative TEM images of CHIR+ cultures treated with 5μM of XL413 hydrochloride or 1μM PD0332991 showing cells with multiple big immature lamellar bodies and multivesicular bodies (representative of 1 experiment).

### NOTCH regulates proximodistal specification

Several mouse genetic models suggest a role of NOTCH signaling in proximodistal specification. Depending on the model, however, conflicting results have been reported (Xu et al., 2012). We therefore used our *in vitro* multilineage differentiation model to investigate whether and how NOTCH signaling affects the balance between proximal and distal cells.

In CHIR− cultures, several NOTCH receptors and targets **(Fig. 8a)** and NOTCH1 intracellular domain (NICD) **(Fig. 8b)** were upregulated compared to CHIR+ cultures, indicating that CHIR inhibits NOTCH signaling or that CHIR withdrawal induced a population that is subject to NOTCH signaling. To examine the role of NOTCH signaling in CHIR− cultures, we added the *γ*-secretase inhibitor, DAPT. RT-qPCR **(Fig. 8c)**, IF **(Fig. 8d)**, and FCM **(Fig. 8e)** (comparative IF for 18 markers shown in **Fig. S1)** showed that inhibiting NOTCH in CHIR− cultures favored generation of ITGA6^+^ITGB4^+^ BC-like cells, which made up 45±10.5% of the population (not shown), as well as ciliated (FOXJ1 and AcTUB) and neuroendocrine (CALCA, UCHL1) fates while profoundly inhibiting club cell (SCGB3A2) and distal fates. Therefore, NOTCH inhibition induced further differentiation of a select set of airway cells at the expense of other fates, indicating a role for NOTCH signaling in proximodistal specification.

**Figure 8.**
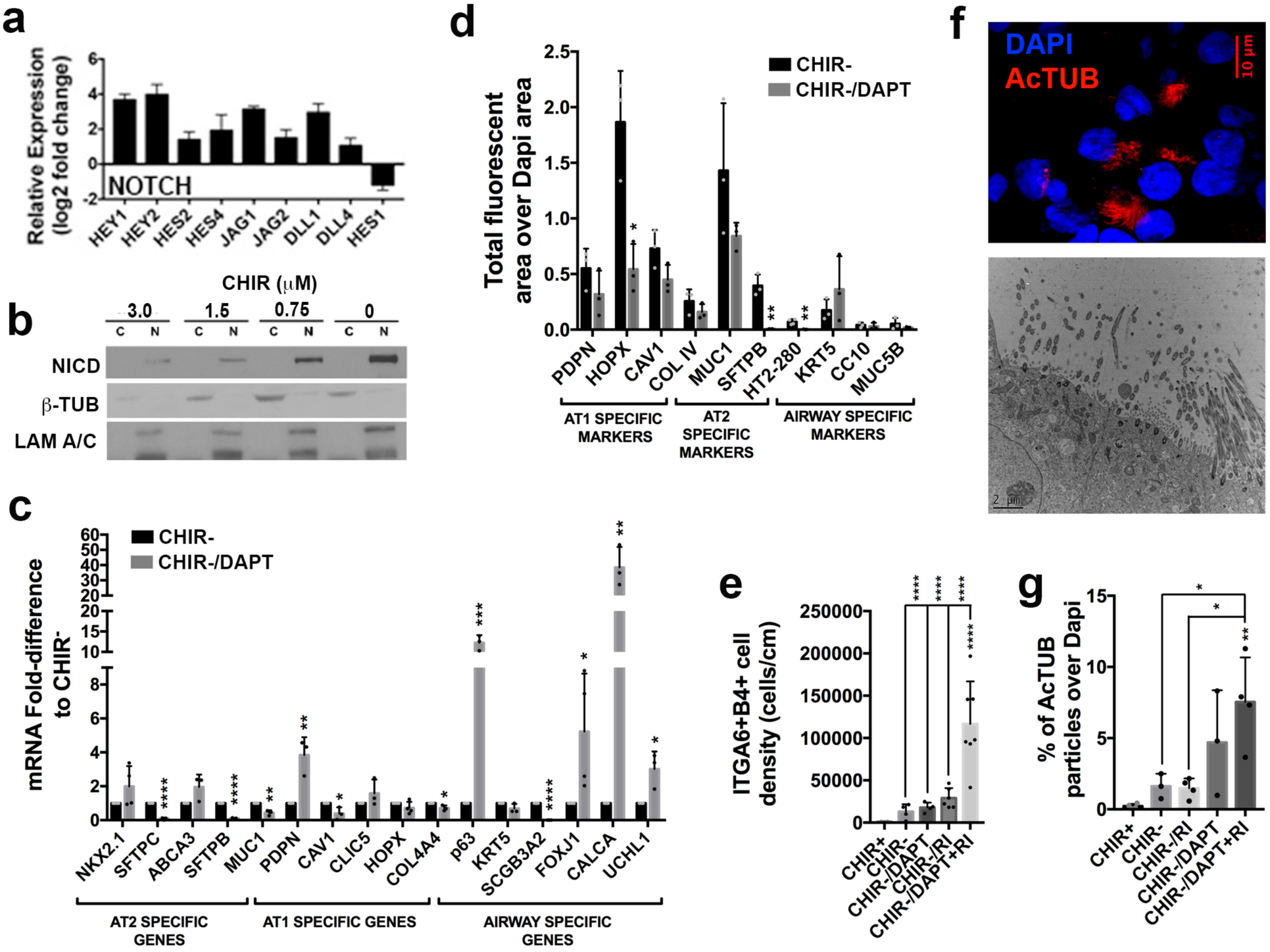
Effect of NOTCH inhibition on differentiation in CHIR− cultures. **(a)** NOTCH-related genes significantly differently expressed by RNAseq in CHIR− compared to CHIR+ cultures (mean± s.d, n=3 independent experiments, all data shown FDR<0.05). **(b)** WB on nuclear and cytoplasmic fractions for NOTCH intracellular domain (NICD) in cultures treated with varying concentration of CHIR (representative of 3 experiments). **(c)** Expression of mRNA for differentiation markers in CHIR− cultures treated with 25μM of DAPT relative to CHIR− cultures (mean± s.d., n=3-4 independent experiments, *p<0.05, **p<0.01, ***p<0.001, ****p<0.0001). **(d)** Relative quantification of IF for non-nuclear protein as determined by the ratio between fluorescent area for a given marker and nuclear DAPI area in CHIR− cultures and CHIR− cultures treated 25μM of DAPT (mean± s.d., n=3-4 independent experiments, *p<0.05, **p<0.01). **(e)** Density of ITGA6^+^ITGB4^+^ cells in CHIR+, CHIR−, and CHIR− cultures treated with 25μM of DAPT and/or 10μM of RI as determined by FCM. (mean± s.d., n=4-7 independent experiments, ****p<0.0001). **(f)** Representative confocal image of AcTUB (representative of 4 independent experiments) and TEM image (representative of 1 experiment) of a cluster of ciliated cells in CHIR− culture supplemented with 25μM of DAPT and 10μM of RI. **(g)** Fraction of AcTUB^+^ clusters relative to DAPI in CHIR+, CHIR−, and CHIR− cultures treated with 25μM of DAPT and/or 10μM of RI as determined by IF. (mean± s.d., n=3-4 independent experiments, *p<0.05, **p<0.01).

As inhibition of Rho-associated kinase (ROCK) facilitates culture of epithelial progenitors (Butler et al., 2016; Mou et al., 2016) and as DAPT promoted select proximal fates, we combined DAPT with the ROCK inhibitor (RI), Y27632. This resulted in a marked synergistic increase in ITGA6^+^ITGB4^+^ (1.17±0.5×10^5^ cells cm^−2^, 66.4±4.04%) (**Fig. 8e**) and P63^+^ cells (57.4±14.46%) (not shown). This combination also further enhanced generation of AcTUB^+^ multiciliated cells (7.5±3.12%) **(Fig. 8g)** with numerous long cilia **(Fig. 8f)** confirmed by TEM **(Fig. 8f)** and by high-speed live imaging **(Suppl. Movie 1,2)**. Taken together, these findings indicate a roadmap for the generation of proximal progenitors and ciliated cells from LPs through removal of CHIR followed by inhibition of both NOTCH and ROCK signaling.

### Reproducibility in iPS lines

To ascertain the general validity of our observations made using RUES2 ESCs, we reproduced key findings in two iPS lines **(Fig. S7)**. In both lines, CHIR removal inhibited cellular expansion **(Fig. S7a,b)** and induced a wide array of markers of differentiating cells **(Fig. S7c,d).** Furthermore, addition of DAPT and RI to CHIR− cultures strongly increased the absolute number **(Fig. S7e)** and the fraction **(Fig. S7f)** of ITGA6^+^ITGB4^+^ cells. Both iPS lines therefore behaved similarly to RUES2 ESCs.

## Discussion

We described here a method to differentiate LPs into the most mature lung and airway epithelial cells reported yet by culture in Col I 3D media in the absence of GSK3i. Neither pseudostratified epithelium, nor NGFR^+^ BCs, nor ATI cells that are morphologically recognizable as such were ever generated *in vitro,* although the lack of expression AGER on ATI suggests that these are still not fully mature. Importantly, in contrast to previous reports (Jacob et al., 2017; McCauley et al., 2017), this approach does not require reporter lines to isolate LPs or to identify specific lineages.

Our observations indicate that GSK3, a kinase that integrates multiple inputs and affects a wide variety of signaling pathways and cellular process in addition to WNT signaling (Patel and Woodgett, 2017), is a pivotal node in the regulation of the differentiation of LPs that appears at least partially independent from its effect on WNT signaling. The generation of embryonic lung progenitors (LPs) from AFE requires WNT signaling *in vitro* (Huang et al., 2014) and *in vivo (Goss et al., 2009)*, which is stimulated *in vitro* by addition of the small molecular GSK3 inhibitor, CHIR99021 (An et al., 2010). Continued presence of CHIR promoted cellular expansion and inhibited multilineage differentiation. Although canonical WNT signaling inhibits GSK3 Ser/Thr kinase activity, thereby stabilizing *β*-catenin (Willert and Nusse, 2012) the effect of CHIR could not be qualitatively recapitulated by a canonical WNT ligand, WNT3a. In fact, canonical WNT ligand promoted a proximal fate, whereas CHIR repressed both proximal and distal maturation. While WNT3A may be less active than CHIR with respect to activation of canonical WNT signaling in human cells (Fuerer and Nusse, 2010), rhWNT3A did induce the WNT target AXIN2 **(Fig. 6f)**, indicating biological activity. Furthermore, as the effect of CHIR was dose-dependent **(Fig. 3b,d)**, a threshold effect of WNT signaling is unlikely. The effect of GSK3 could therefore not be assigned to a single signaling pathway. Part of the effect of CHIR may involve enhancement of cell cycling, one of the integrated outputs of GSK3, in particular for distal fates. This is consistent with the notion that GSK3 is a central node in cellular physiology that integrates multiple inputs and regulates a broad range of downstream processes, including cell cycle (Patel and Woodgett, 2017). Similar observation were made in radial glia during brain development, where cell cycle inhibition favored differentiation at the expense of self-renewal (Calegari et al., 2005; Calegari and Huttner, 2003; Kim et al., 2009; Lange et al., 2009; Roccio et al., 2013). It is also clear however that mere cell cycle inhibition does not reproduce all effects of CHIR withdrawal, as CHIR withdrawal induced much more pronounced proximal differentiation, as well as morphologically more advanced ATII maturation.

Our findings appear at odds with reports showing that timed removal of CHIR from LPs promoted a proximal fate (McCauley et al., 2017), that its continued presence drove an ATII fate (Jacob et al., 2017) and that these effects were explained by attenuation or agonism of WNT signaling, respectively (Jacob et al., 2017; McCauley et al., 2017). In our Col I 3D model, WNT3a in fact induced a proximal and club cell fate. This is the opposite of the findings of McCauley et al. and Jacob et al. (Jacob et al., 2017; McCauley et al., 2017), who used Matrigel to read out terminal cell fate. Although Col I is more permissive than Matrigel for the development of a broad array of lung and airway lineages, we obtained qualitatively similar data in Matrigel. A difference in the 3D media used can therefore not fully explain this discrepancy. McCauley et al. withdrew CHIR as early as d15, at the early LP stage. We observed however that at that stage, the cells were not yet fully competent to differentiate in 3D culture after CHIR withdrawal. We furthermore also found that CHIR withdrawal at d35 had a similar effect as withdrawal at d25. Our studies therefore indicate that this discrepancy is not explained by different timing of CHIR withdrawal. McCauley et al. (McCauley et al., 2017) and Jacob et al. (Jacob et al., 2017) used reporter lines and cell sorting to isolate LPs from early-stage, low-purity cultures and withdrew CHIR in 2D culture prior to reading out proximal (McCauley et al., 2017) or distal fate (Jacob et al., 2017) in Matrigel spheroid cultures in conditions that are selective for one or the other fate. In contrast, we did not perform cell sorting and withdrew CHIR after plating in Col I. This may be important as we observed that during prolonged 2D culture detachment of cells occurs, which may, depending on culture conditions, affect some cell types more than others (Huang et al., 2015). Furthermore, to identify distal potential they first isolated SFTPC-reporter positive cells, which are already committed to the ATII lineage. It therefore appears logical that only ATII cells are detected in subsequent 3D cultures. Interestingly, however, they do report that withdrawal of CHIR in Matrigel cultures of ATII progenitors induced maturation, a finding that may be consistent with our observation that CHIR withdrawal induces multilineage differentiation, including differentiation of morphologically mature ATII cells. We also note that we have previously shown that even immature ATII cells take-up and recycle SFTPB (Chen et al., 2017; Huang et al., 2014). Surfactant recycling is therefore a functional characteristic that is likely acquired relatively early in ATII differentiation and is not an indicator of maturity.

We next performed studies to examine whether this model can be used to gain deeper insight into proximodistal specification. A canonical WNT ligand stimulated club cell generation, while NOTCH inhibition promoted proximal, neuroendocrine and ciliated fates at the expense of distal fates. Although the role of NOTCH signaling in distal lung development *in vivo* is unclear (Xu et al., 2012), our observations on its effect on the induction of specific proximal fates is consistent with mouse genetic models (Mori et al., 2015; Rock et al., 2011). NOTCH was furthermore synergistic with ROCK inhibition in this respect. This was surprising, as it has been suggested that RI conditionally immortalizes a variety of epithelial progenitors by suppressing non-canonical NOTCH signaling pathways (Yugawa et al., 2013).

Collectively, our observations show that this model provides novel mechanistic insight into human lung development that complement mouse genetic models, and begins to provide a roadmap for the generation of specific lung and airway lineages.

## Acknowledgements

This work was supported by grant NIH HL120046-01 (HWS) and 1U01HL134760-01 as well as funding from the Thomas R Kully IPF Research Fund (HWS). A.L.R.T.C. is recipient of a PhD fellowship (PD/BD/52320/2013) in the context of the University of Minho MD/PhD Program funded by the Portuguese Science Foundation (FCT). Work in RBV’s lab was supported by an NIH HD40182 to RBV and an AHA/ASA postdoctoral fellowship to TJD. We thank NYULMC OCS Microscopy core for their assistance with transmission electron microscopy. We thank NYULMC DART Microscopy Lab, Joseph Sall, Jessica Shivas Ph.D. and Albert E. Ayoub Ph.D for their assistance in the light sheet microscopy work. We thank Dr. Matthew Bacchetta for providing human adult lung samples. Flow cytometry was performed in the CCTI Flow Cytometry Core, supported in part by the Office of the Director, National Institutes of Health under awards S10RR027050 and S10OD020056.

## Authors Contributions

ALRTC co-designed and performed most experiments, contributed to the concept, and co-wrote the manuscript with HWS. YWC and HYL assisted ALRTC in developing and analyzing Col I cultures. AS assisted with all bioinformatics. TJD and RBV provided expertise, support and instrumentarium in the analysis and imaging of ciliated cells. JCP provided mentoring and assistance to ALRTC. HWS provided concept and guidance, co-designed experiments and wrote the manuscript.

## Competing Financial Interests

The authors have no competing financial interests.

## Methods

### hPSC Maintenance

RUES 2 (Rockfeller University Embryonic Stem Cell Line 2, NIH approval number NIHhESC-09-0013, registration number 0013; passage 17-28), Sendai Virus (sv) and modified mRNA generated human dermal fibroblasts iPSC lines (from healthy fibroblasts, purchased from Mount Sinai Stem Cell Core facility, passage 17-26) were cultured on mouse embryonic fibroblasts (GlobalStem, Rockville, MD) plated at 17,000-20,000 cells cm^−2^. hPSC maintenance media consisted of DMEM/F12 (Cellgro, Manassas, VA) 20% Knockout Serum Replacement (Gibco, Life Technologies, Grand Island, NY), 0.1mM β-mercaptoethanol (Sigma-Aldrich, St. Louis, MO) 1 % GlutaMax (Gibco), 1% Non-essential amino acids (Gibco), 0.2% Primocin (InvivoGen, San Diego, CA) and 20ng ml^−1^ FGF-2 (R&D System, St. Louis, MO). Media was changed daily and cells were passaged every 4 to 5 days using Accutase/EDTA (Innovative Cell Technologies, San Diego, CA) at 1:24 dilution. Cells were maintained in an undifferentiated state in a humidified 5% CO_2_ atmosphere at 37°C.

### Serum Free Differentiation (SFD) Media

All differentiations were carried in serum free media consisting of IMDM/Ham’s F12 (3:1) (Cellgro), N2 and B27 supplements (Gibco), 1% GlutaMax, 1% penicillin-streptomycin (Cellgro), 0.05% bovine serum albumin (BSA) (Gibco). 50μg ml^−1^ of ascorbic acid (Sigma-Aldrich), 0.4μM monothioglycerol (Sigma-Aldrich) and the indicated growth factors were added fresh.

### Definitive Endoderm Induction (days 1-4 differentiation protocol (dp))

hPSCs were dissociated with Accutase/EDTA for 3min at 37°C to small 3-10 cell clumps and plated to low attachment 6-well plates at 120,000-150,000 cells cm^−2^ to form embryoid bodies (EBs) in SFD media supplemented with 100ng mL^−1^ Activin A (R&D System), 10μM of rock inhibitor (RI) Y-27632 (Tocris, Bristol, BS, UK), 0.5ng mL^−1^ BMP4 (R&D System) and 2.5ng mL^−1^ FGF2 (R&D System) for 72h at 37°C in 5% CO2 5% O2 95% N2 atmosphere. Complete media changes were performed every 24h. Endoderm induction efficiency was determined by the percentage of double positive cells for CXCR4 and cKIT by FCM as previously described^1^,^2^ after dissociation of EBs with 0.05% trypsin/EDTA (Gibco). Typically experiments with ≥98% CXCR4^+^cKit^+^ cells for RUES2 and ≥ 90% CXCR4+cKit+ cells for sv and mRNA iPSC were carried further.

### Anterior Foregut Endoderm Induction (days 4-6 dp)

Anterior foregut endoderm induction (AFE) was carried out as previously described^1^,^2^. Briefly EBs were dissociated to single cells with 0.05% trypsin/EDTA for 5min and plated at 50,000-75,000 cells cm^−2^ in 0.33% fibronectin (R&D System) coated 24-well plates in SFD media supplemented with 2μM Dorsomorphin (Tocris) and 10μM SB431542 (Tocris) for the first 24h followed by SFD supplemented with 1μM IWP2 (Tocris) and 10μM SB431542 for another 24h. Differentiation was carried at 37°C in 5% CO_2_ 5% O_2_ 95% N_2_ atmosphere.

### Lung Progenitor Specification (days 6-15 dp)

Lung Progenitors (LPs) specification was carried out as previously described^1,2^. Briefly a media switch was performed following the 48h of AFE induction to SFD plus the lung factors 3μM CHIR99021 (Tocris), 10ng mL^−1^ FGF7 (R&D System), 10ng mL^−1^ FGF10 (R&D System), 10ng mL^−1^ BMP4 and 50nM all-trans retinoic acid (R&D System). Complete media changes were performed every 48h and cells were maintained at 37°C in 5% CO_2_ 5% O_2_ 95% N_2_ atmosphere for the first 2 to 3 days in lung factors media and then switched to 5% CO_2_ atmosphere at 37°C.

### Expansion of LPs (days 15-25 dp)

At d15 of the differentiation protocol cells were briefly trypsinized for 1min at 37°C^1^,^2^ and replated into growth factor reduced matrigel (Corning, Corning, NY) (1:100 dilution) 24-well coated plates at 1:2 splitting ratio. LPs were expanded in SFD plus 3μM CHIR99021, 10ng mL^−1^ FGF7 and 10ng mL^−1^ FGF10, with media changes every 48h and in 5% CO_2_ atmosphere at 37°C. At d25 experiments were accessed for LPs generation efficiency by immunofluorescence staining for Nkx2.1, Sox2 and Foxa1 as previously described^1,2^. Typically experiments with ≥90% Nkx2.1^+^Sox2^+^Foxa1^+^ cells for RUES2 and ≥ 80% Nkx2.1^+^Sox2^+^Foxa1^+^ for sv and mRNA iPSC were carried further.

### LP Maturation in Collagen I gels (days 25-50 dp)

At d25 of the differentiation protocol cells were briefly trypsinized for 1min at 37°C. Cells were scraped off the plate and cell clumps collected and re-suspended in a 4.5 mg mL^−1^ rat tail collagen I (Col I) (Trevigen, Gaithersburg, MD) solution prepared according to manufacturer’s gelation protocols. An equivalent of 100,000 cells cm^−2^ in cell clumps were plated in 24 or 48-well plates and placed at 37°C 10-15min for gelation. Media was added on top of the gels and changed every 48h. Maturation media consisted of SFD plus 10ng mL^−1^ FGF7, 10ng mL^−1^ FGF10, 50ng mL^−1^ Dexamethasone (Tocris), 0.1mM 8-bromo-cAMP (Tocris) and 0.1mM IBMX (Tocris) (CHIR−). When indicated CHIR99021 was included in the maturation media (CHIR+) (3μM). For some experimental conditions other factors or small molecules were added to the media: 25μM DAPT (Tocris), 10 μM Y-27632 (RI), 100ng mL^−1^ WNT3A (R&D System) 1μM PD0332991 (Tocris) and 5μM XL413 hydrochloride (Tocris). Cells were maintained in a 5% CO2 atmosphere at 37°C.

### Timed CHIR withdrawal

For experiments in which CHIR was retracted at d15, d15 LPs were briefly trypsinized for 1min at 37°C^1,2^. Cells were scraped off the plate and cell clumps collected and re-suspended in 4.5 mg mL^−1^ rat tail Col I solution at an equivalent of 100,000 cells cm^−2^ in cell clumps plated in 48-well plates and placed at 37°C 10-15min for gelation. Cultures were maintained in CHIR+ or CHIR− differentiation media as described in the previous section until d50. For experiments in which CHIR was retracted at d35, 3D cultures were established as described above at d25 and maintained in CHIR+ media until d35, after which CHIR was retracted and cultures maintained in CHIR− maturation media until d50.

### LPs Maturation in Matrigel 3D Media

d25 LPs were briefly trypsinized and scraped off the tissue culture plate as described above. An equivalent of 100,000 cells cm^−2^ in cell clumps were suspended in Matrigel 3D media and plated in 48-well plates. After polymerization at 37°C cultures were maintained in CHIR+ or CHIR− differentiation media as described above until d50. For cultures maintained in CHIR− FGF2 maturation media, SFD was supplemented with 250ng mL^−1^ FGF2, 100ng mL^−1^ FGF10, 50ng mL^−1^ Dexamethasone, 0.1mM 8-bromo-cAMP and 0.1mM IBMX.

### Human Lung Tissue

Adult lung samples were obtained from lungs rejected for transplantation procured from the LiveOnNY (Live On New York) organ procurement organization, under a protocol approved by the Institutional Review Board at Columbia University.

### Immunofluorescence

d50 Col I gels, Matrigel 3D cultures and human lung samples were embedded in OCT and sectioned into 6μm slices placed on microscopic slides. Slices were briefly fixed with 95% ethanol for 1min at room temperature (RT) followed by 4% paraformaldehyde 15min RT. After washing twice with PBS slices were permeabilized with 0.25% triton in blocking buffer (5% donkey serum (EMD Millipore, Darmstadt, Germany) in PBS) for 10min RT and blocked for 1h RT in blocking buffer. Primary antibodies diluted in blocking buffer were incubated at 4°C overnight. In the following day, slices were washed twice with PBS and secondary antibodies (Jackson ImmunoResearch, West Grove,PA) (Invitrogen) diluted in blocking buffer were incubated for 2h RT followed by a 5 min incubation with Dapi in PBS and 2 washes with PBS. Slides were mounted with ProLong Gold antifade reagent (Invitrogen (Thermo Fisher Scientific), Waltham, MA) sealed and preserved in dark at 4°C. A complete list of antibodies and dilutions used can be found in **Supplementary Table 2**.

ALI cultures were fixed with 4% paraformaldehyde 15min RT prior to embedding in OCT and sectioned into slides. Sorted ITGA6^+^ITGB4^+^ cells were plated into Col I coated glass bottom plates, spun down at 70g and left 1h at 37°C to adhere before being fixed and stained as described above.

Samples were imaged using a motorized Leica DMI6000 B or DMi8 (Leica-Microsystems, Buffalo Grove, IL) inverted microscopes and processed using ImageJ software. For high magnification images we used IX83 Andor Revolution XD Spinning Disk Confocal System with a 100x oil objective (NA 1.49) and a 2x magnifier coupled to an iXon Ultra 888 EMCCD Camera.

### Immunofluorescence quantification

For each marker tile scans corresponding to the whole cross section area were acquired and quantified using ImageJ software. For transcription factors, images were converted to 8-bit and threshold values were determined to cover nuclear area. Number of positive particles (positive nuclei) was determined by ‘Analyze Particles’ function of ImageJ. Percentage of positive cells was calculated by dividing the number of positive particles by the total number of dapi stained particles (total number of nuclei). For cytoplasmic and membrane markers, total positive (fluorescent) area was calculated by converting the image to 8-bit and thresholding it to the value that best covered the stained area. Fluorescent area was determined by ‘Analyze Particles’ function. Total fluorescent area was then normalized to total dapi stained area.

### Quantitative real-time PCR

d50 cultures in Col I gels were digested with 150U mL^−1^ collagenase type I in IMDM for 45min at 37°C. Cells were collected and total RNA was extracted using RLT buffer and RNeasy Micro Kit (Qiagen, Valencia, CA). RNA concentration was measured with NanoDrop 2000 fluorospectrometer (Thermo Fisher Scientific). cDNA was generated by reverse transcription of 1μg of total RNA with random hexamers and Superscript III (Invitrogen) following manufacturer’s instructions. Real-time quantitative PCR was performed using ABI Power SYBR green PCR Master Mix (Applied Biosystems (Thermo Fisher Scientific)) on ABI vii7A Thermocycler (Applied Biosystems) with the following amplification conditions: 50°C for 2min and 95°C for 10min followed by 40 cycles of 95°C for 15s and 60°C for 1min plus dissociation/melt curves. Real time quantitative PCR was performed on three biological samples from each experiment with technical triplicates for each sample. cDNA input per reaction was 5ng. Analysis was performed using the standard curve method - for each gene absolute quantification was obtained using a standard curve of serial diluted genomic DNA and normalized to housekeeping gene TBP (Tata Box Binding protein), then fold-differences were calculated to designated calibrator sample (CHIR^+^ or CHIR^−^ conditions). Used primer sequences can be found in **Supplementary Table 3**.

### Flow Cytometry

d50 cultures in Col I gels were digested with 150U mL^−1^ collagenase type I in IMDM for 45min at 37°C. Cell colonies were collected and further dissociated with pre-warm 0.05% Trypsin/EDTA for 5min to single cells. Cells were stained at 4°C for 45min in PBS with 0.2% BSA and 2mM EDTA with (1) anti-EpCAM PerCP (Invitrogen), anti-ITGA6 alexa fluor 647 and anti-ITGB4 PE (Biolegend, San Diego, CA); (2) anti-EpCAM PerCP and anti-NGFR APC (Biolegend) or (3) anti-EpCAM and unconjugated HT2-280 antibody followed by anti-mouse IgM Alexa Fluor 647. Stained cells were analyzed on BD LSRII and BD Fortessa Analyzers (BD Bioscience, San Jose, CA). Results were analyzed in Flowjo v10.2 software. Analysis was gated on live, doublet-excluded, EPCAM^+^ cells.

### Population Isolation by FACS sorting

d50 cells were dissociated and stained for EPCAM, ITGA6 and ITGB4 as described above. Populations were sorted using a BD Influx cell sorter (BD Bioscience) into complete media.

### NGFR^+^ Basal cell expansion and Air-Liquid Interface cultures

COL 1 gels were digested with collagenase type 1 and cell clumps trypsinized to single cells as described above. Single cells were stained with anti-human EpCAM-PerCP, anti-human ITGB4-PE and anti-human NGFR-APC for 45min at 4C in FACS buffer. EpCAM^+^ITGB4^+^NGFR^+^ basal cells were sorted using a BD Influx cell sorter (BD Bioscience) into epithelial media^4^. Sorted cells were seeded onto irradiated 3T3-J2 feeders and maintained in epithelial media^4^ over 3 passages. Passage 2 cells were used for air-liquid interface cultures. 200.000 cells were seeded onto matrigel coated (1:30) 0.33cm^2^ transwells and kept in epithelial media^4^ on the upper and lower chamber for the first 48h, after which ALI was induced and cells maintained in PneumaCult-ALI media (Stem Cell Technologies, Cambridge, MA).

### Western Blot

d50 cultures in Col I gels were digested with 150U mL^−1^ collagenase type I in IMDM for 45min at 37°C. Cell colonies were collected and further dissociated with pre-warm 0.05% Trypsin/EDTA for 3-5min to small cell clumps (≤10cells). Nuclear/cytosol cell fractionations were performed by lysing the cell pellets with a 2x volume of 10mM HEPES pH8.0, 1.5mM MgCl_2_ solution with 1x protease inhibitors (Roche, Basel, Switzerland) and 1/10 volume of 3% NP-40 for 10min on ice. Plasma membrane lysis was verified by trypan blue staining. Lysate was spun at 21,000g for 10min at 4°C and the supernatant (cytoplasmic fraction) was collected. Remaining pellet was re-suspended in 1x volume of 20mM HEPES pH8.0, 25% glycerol, 420mM NaCl, 1.5mM MgCl_2_, 0.2mM EDTA solution with 1x protease inhibitors, incubated at 4°C for 1h with rotation and spun at 21,000g for 20min at 4°C. Supernatant (nuclear fraction) was collected and diluted with 1x volume of 1.5mM MgCL2, 0.6% NP40 and 0.2mM EDTA nuclear diluent solution. For each sample 20μg of protein corresponding to nuclear and cytosolic fractions were denaturated in 6x loading buffer at 95°C for 5min and loaded onto separate lanes of 4-12% Bis-Tris SDS-PAGE gradient Gels (Invitrogen). Gels were transferred into a 0.22μm nitrocellulose membrane and stained with Ruby Red (Molecular probes, Carlsbad, CA) to confirm transfer. Membranes were blocked with 5% non-fat milk in TBS-Tween 0.05% and incubated with anti-cleaved NOTCH1, anti-β-catenin, anti-phospho-GSK3β (Ser9), anti-Lamin A/C, anti-β Tubulin antibodies overnight at 4°C with agitation. Membranes were washed, incubated with the appropriate HRP-conjugated secondary antibodies for 2h RT with agitation, washed again and exposed to X-ray film (Denville, Holliston, MA) after incubation with Pierce ECL Western Blotting Substrate or SuperSignal West Femto ECL reagent (Thermo Fisher Scientific). A complete list of antibodies and dilutions used can be found in **Supplementary Table 2**.

### RNA sequencing

Total RNA from d25 and d50 (after Col I digestion as described above) cultures was purified with RNeasy Micro Kit. Agilant microfluidic RNA 6000 Nano Chip kit (Agilent Technologies, Santa Clara, CA) and 2100 Bioanalyser (Agilent Technologies) were used to determine RNA concentration an integrity number (RIN). All sequenced samples had RIN ≥9.5. Poly-A pull-down was used to enrich mRNAs from total RNA samples. Library preparation was performed with Illumina TruSeq RNA prep kit (Illumina, San Diego,CA). Libraries were then sequenced using Illumina HiSeq2500 at Columbia Genome Center. Samples were multiplexed in each lane to yield targeted number of single-end 100bp reads for each sample, as a fraction of 280-400 million reads for the whole lane. RTA (Illumina) was used for base calling and bcl2fastq2 (version 2.17) for converting BCL to fastq format, coupled with adaptor trimming. Reads were mapped to human genome (NCBI/build37.2) using STAR(2.5.2b) and featureCounts (v1.5.0-p3). Differentially expressed genes under various conditions were tested using DEseq, an R package based on a negative binomial distribution that models the number reads from RNA-seq experiments and test for differential expression.

### Single Cell RNA sequencing

d50 cultures in Col 1 were digested with collagenase type I as described aboved, followed by trypsinization with pre-warmed 0.05% Trypsin/EDTA until cultures were dissociated to single cells. The Chromium™ Single Cell 3’ Library & Gel Bead Kit v2, 16 rxns PN-120237 kit was used according to manufacturer’s instructions. 5000 cells/library were targeted. 12 cycles were used for cDNA Amplification and for the Sample Index PCR. Libraries were pooled and sequenced to a depth of ∼350M reads on an Illumina HiSeq 4000. 10x genomics’ cellranger pipeline v2.0.0 was used to process the data and the reference genome was GRCh38.

### KeyGenes analysis

Identity of d25 and d50 cultures were predicted by KeyGenes algorithm^4^ from its transcriptional profile based on next generation sequencing data of human fetal tissues from the first and second trimester of development and adult tissues.

### Transmission Electron Microscopy

TEM was performed at NYULMC Microscopy Core Laboratory. Col I gels were fixed with 2.5% glutaraldehyde and 2% paraformaldehyde in 0.1M sodium cacodylate buffer (pH7.2) for 2 hours and post-fixed with 1% osmium tetroxide for 1.5 hours at room temperature, then processed in a standard manner and embedded in EMbed 812 (Electron Microscopy Sciences, Hatfield, PA). Semi-thin sections were cut at 1 μm and stained with 1% Toluidine Blue to evaluate the quality of preservation and find the area of interest. Ultrathin sections (60 nm) were cut, mounted on copper grids and stained with uranyl acetate and lead citrate by standard methods. Stained grids were examined under Philips CM-12 electron microscope and photographed with a Gatan (4k ×2.7k) digital camera.

### Light Sheet Microscopy

d50 cultures within the Col 1 gels were fixed overnight with 4% PFA, washed three times with PBS for 10min and permeabilized with 0.25% triton in blocking buffer (5% donkey serum in PBS) for 4h RT with rotation. The gels were blocked with blocking buffer for 4h RT with rotation and then left incubating with primary antibodies overnight at 4C with rotation. In the following day the gels were washed three times with PBS for 10min with rotation, incubated with the appropriate secondary antibodies for 4h RT with rotation followed by 10min incubation with DAPI in PBS and two 10min washes with PBS at RT with rotation. 3D structures within the gel were dissected, mounted on agarose and imaged with the SP8-DLS system on a DMi8 inverted microscope, 2.5x illumination objectives with the 5mm TwinFlect mirror fitted on the 10x detection objective (0.3 NA W DLS) (Leica-Microsystems, Buffalo Grove, IL) or the Zeiss Lightsheet Z.1 running ZEN 2014 SP1 (black edition) version 9,2, detection objective EC Plan-Neofluar 5x/0.16NA (Carl Zeiss Microscopy, Thornwood, NY). Acquired images were then resized and 3D renderings were obtained using the 3D project function of ImageJ Software.

### Live Imaging of d50 cultures

d50 cultures in Col I gels were digested with 150U mL^−1^ collagenase type I (Gibco) in IMDM for 45min at 37°C. Cell colonies were collected and further dissociated with pre-warm 0.05% Trypsin/EDTA for 2-3min to small sized cell clumps. Cells were re-suspended in SFD maturation media and plated in 50mm glass-bottom dishes. Live imaging was performed using an IX83 Andor Revolution XD Spinning Disk Confocal System with an environmental chamber at 37°C, a 60x silicone oil objective (NA 1.30). Image acquisition was carried out using a Zyla 5.5. sCMOS camera (2048 × 2048 px) and the imaging software Metamorph.

### Statistics and reproducibility

For statistical analysis between two groups unpaired two-tailed Student’s *t*-test was used. For multiple group comparison one-way ANOVA was performed followed by Dunnett or Tukey multiple comparison tests. Results are displayed as mean ± s.d. with *p* values <0.05 considered statistically significant. N-values refer to biologically independent replicates. Grubbs and ROUT tests were used to exclude outliers.

### Data availability

We have uploaded the RNA sequencing and single cell RNA sequencing results to NCBI GEO database with the following accession number GSE101558. All data supporting the findings in this manuscript are available as **Supplementary Table 4.** All other data supporting the findings in this manuscript are available upon reasonable request from the corresponding author.

